# Evaluation of real-time PCR performance for detecting *Diaporthe destruens* in sweet potatoes

**DOI:** 10.1101/2025.03.03.641124

**Authors:** Takashi Fujikawa, Yasuhiro Inoue, Yoshifumi Tsukiori, Hiroyoshi Inoue, Kazuki Fujiwara

**Affiliations:** Institute for Plant Protection, National Agriculture and Food Research Organization (NARO), 2-1-18 Kannondai, Tsukuba, Ibaraki 305-8666, Japan; Kyushu Okinawa Agricultural Research Center, NARO, 2421 Suya, Koshi, Kumamoto 861-1192, Japan; Faculty of Agriculture, Meijo University, 501 Siogamaguchi 1, Tenpaku-ku, Nagoya 468-8502, Aichi, Japan

**Keywords:** Foot rot disease, real-time PCR, detection performance, a receiver operating characteristic (ROC) curve

## Abstract

Sweet potato foot rot disease caused by *Diaporthe destruens* is a major threat to sweet potato production. Rapid and accurate detection in advance is essential to secure healthy plants or to implement effective control. In this study, we evaluated the detection performance of a real-time PCR method previously developed by our research group. The evaluation was conducted using sweet potato samples collected in the field, including those naturally infected with *D. destruens*. Based on the real-time PCR results and the actual occurrence of the disease, a receiver operating characteristic (ROC) curve was generated, and the area under the curve (AUC) was calculated. The AUC values ranged from 0.7041571 to 0.9204286, confirming that this real-time PCR method is a well-balanced detection method for both tubers and stems. Furthermore, to minimize the occurrence of false negatives, the cycle threshold (Ct) cutoff was set at 35, and detection performance (sensitivity, specificity, accuracy, etc.) was analyzed. Then, by considering abnormal melting temperature (Tm) values as negative, high accuracy was achieved, with values of 0.871212 for tubers and 0.715447 for stems. This study not only evaluated the detection performance of the real-time PCR for foot rot pathogens but also contributes to providing information on the methodology and significance of performance evaluation in various real-time PCR genetic detection methods.

## Introduction

Sweet potato foot rot, caused by a fungus *Diaporthe destruens* (Harter) Hirooka, Minosh. & Rossman, is a serious disease affecting *Ipomoea batatas* (L.) Lam. (Gai et al. 2016; Maeda et al. 2022). Infected plants develop blackening and wilting of the stems (foot rot), which lead finally to plant death. Additionally, infected root tubers often rot during storage. In severely affected fields, black, hardened stem lesions and rotting tubers in the soil are commonly observed (Gai et al. 2016; Kobayashi 2019; Maeda et al. 2022; Nishioka et al. 2021). First identified in the United States in 1912, the disease has since found in Cuba, the Caribbean, Brazil, Argentina, eastern Africa, New Zealand, and other regions (Fujiwara et al. 2021; Gai et al. 2016; Harter 1913). More recently, its presence has been confirmed in East Asia, including Japan, Taiwan, China, and Korea, where it has become a major challenge for sweet potato cultivation (Gai et al. 2016; Huang et al. 2012; Paul et al. 2019).

The primary source of transmission of this pathogen is through planting infected seedlings (either stem cuttings or seed tubers) for propagation (Usui and Kushima 2020). In addition, if infected plants are left in the field, the pathogen will grow on the residue and wait in the soil for the next transmission (NARO et al. 2023). More, the infection will spread when healthy seedlings or tubers come into contact with infected plants (NARO et al. 2023). For this reason, control and prevention of the disease are important both in the field and in seedling production, and rapid and accurate diagnosis of infected plants is essential prior to that. Thus, our group developed a genetic detection method using real-time PCR that can identify foot rot pathogen contamination without relying on visual inspection (Fujiwara et al. 2021; Tsukiori et al. 2023). This method was developed to identify the pathogens of foot rot disease (*D. destruens*) and its similar disease, dry rot disease (*D. batatas*). It is confirmed that this method can detect the pathogens with high sensitivity (Fujiwara et al. 2021). Also, it is robust because it does not misdiagnose other fungal diseases of sweet potatoes (Tsukiori et al. 2023). In Japan, the diagnosis of foot rot disease is a high priority in sweet potato production at present, and this method is becoming more well-known as a diagnostic method for foot rot disease.

While real-time PCR is expected to be rapid and accurate, confusion often arises as to how to interpret the results. Particularly regarding the setting of cycle threshold (Ct) value that distinguish between positive and negative results, and the decision whether to interpret results based on temperature of melting (Tm) value (in the case of intercalator assays such as SYBR Green), which appears to be analyst dependent. In the real-time PCR method of Fujiwara et al. (2021), it has been considered appropriate to exclude Ct values that are easily appeared false positives and abnormal Tm values due to non-specific amplification products, and to judge a result as positive when the Tm value is normal (derived from specific amplification products) and the Ct value is less than 35 (Tsukiori et al. 2023).

Here, we had access to naturally contaminated tubers and seedlings, which were collected from different fields in areas where foot rot was widespread at the time of the start of the study due to insufficient control measures. Using these contaminated plants, the ends of plants (the necks and the tails in the case of tubers, and the basal end in the case of seedlings) were used for the real-time PCR to examine if they were contaminated with foot rot pathogen, and the remaining main parts were planted in a field to examine if foot rot disease developed thereafter, so that the relationship between the real-time PCR results and the actual onset of the disease was clarified. In this study, to assist analysts in determining the Ct value for foot rot diagnosis using this real-time PCR method, the receiver operating characteristic (ROC) curves were generated and the detection performance was analyzed.

## Materials and Methods

### Collection of sweet potato plants

At the start of this study, around 2021, we collected tubers and stems of sweet potatoes (The varieties were mostly “Kogane Sengan”) from fields in various locations where foot rot disease was widespread due to insufficient control measures. We assumed that there was a lot of these plants were naturally infected with the foot rot pathogen. Both ends of the tubers (the neck and the tail) (ca. 1-2cm^3^) and the basal ends of the stems (ca. 1.5-cm length) were cut off, these fragments were used as samples for real-time PCR. The remaining bodies of tubers were divided into two groups, those at the neck and those at the tail. These divided tubers and the remaining stems were transplanted to the sterilized fields and grown for approximately three months to confirm whether the plants were diseased.

### DNA extraction

To generate ROC curves and analyze the detection performance of real-time PCR, DNA extraction was performed following the method described by Fujiwara et al. (2021). Briefly, before DNA extraction, each plant sample was immersed in AP1 buffer from the DNeasy Plant Mini Kit (Qiagen, Hilden, Germany) and ground using a pestle and mortar. DNA was then extracted using the DNeasy Plant Mini Kit according to the manufacturer’s instructions.

### Real-time PCR

Real-time PCR was performed with slight modifications to the method described by Fujiwara et al. (2021) and Tsukiori et al. (2023) as follows. A QuantStudio 3 real-time PCR system (Thermo Fisher Scientific Inc., MA) with KOD SYBR qPCR Mix (Toyobo Co., Ltd., Osaka, Japan) was used according to the manufacturer’s protocol.

The 20-μl reaction mixtures contained 10 μl of qPCR Mix, 0.04 μl of 50× ROX reference dye, 0.5 μl of the respective 10 μM primer sets (forward and reverse) with 2 μl of template DNA (10-fold diluted extracted DNA).

For detection of foot rot disease pathogen, *D. destruens*, the forward primer DdITS-F (5’-GTTTTTATAGTGTATCTCTGAGC-3’) and the reverse primer Dd ITS-R (5’-GGCCTGCCCCCTTAAAAA-3’) (Fujiwara et al. 2021), and for detection of sweet potato internal control, the forward primer Ipo psaB-F (5’-TGTGAAACGTTACCCTGCCA-3’) and the reverse primer Ipo psaB-R (5’-GGACCCGGAGACTTTTTGGT-3’) (Tsukiori et al. 2023) were used respectively. The PCR protocol consisted of initial denaturation at 94 □ for 1 min, and 40 cycles of denaturation at 96 □ for 30 s, annealing at 55 □ for 30 s, and extension at 72 □ for 30 s. Further denaturation at 96 □ for 15 s, holding at 55 □ for 1 min, and heating from 55 to 96 □ for15 s were carried out for melting curve analysis. In this study, the appropriate Tm value was approximately 84°C, which was the same or nearly the same as the Tm value obtained from real-time PCR of diseased plants infected with *D. destruens*, which had been prepared in advance as a positive control, and any Tm value greater or smaller than this was considered abnormal.

### Generating the receiver operating characteristic (ROC) curves

To estimate the performance of real-time PCR assays, the receiver operating characteristic (ROC) curves were generated and analyzed. The ROC curve is a popular graphical method of displaying the discriminatory accuracy of a marker (diagnostic test) for distinguishing between two populations (e.g., healthy and diseased) (Baker 2003; Fluss et al. 2005; Greiner et al. 2000; Hoo et al. 2017; Leach et al. 2018; Nutz et al. 2011). We used 132 tubers collected from actual fields, including those with foot rot, and performed real-time PCR assay using DNA from the neck or the tail sections of tubers. And we visually inspected plants grown from the remaining bodies of tubers for the presence of foot rot. Similarly, we used 163 stems collected from actual fields. We performed real-time PCR assay using DNA from the ends of stems, and visually inspected plants grown from the remaining seedlings for the presence of foot rot.

For each sample, the threshold of cycles (Ct) and the melting temperature (Tm) values obtained by real-time PCR assay, as well as the presence or absence of disease symptoms on the grown plants, were tabled. We also created a table excluding samples with Tm values different from the Tm value obtained by correctly amplifying the target as “outliers.” In real-time PCR assay, the Ct value increases as the original amount of DNA to be detected decreases, the sign of the Ct value is made negative to adjust the correlation between the Ct value and the detection performance to a positive in this table.

In addition, diseased plant samples are represented by a binary value of “1” and healthy plant samples are represented by a binary value of “0.” Based on these tables, we generated the ROC curves with the false positive rate on the horizontal axis and the true positive rate on the vertical axis. Generation of the ROC curve and calculation of the area under the ROC curve (AUC) were performed using the ROCR package in R4.2.1 (e.g., Suppl. Text 1).

### Analytical sensitivity, specificity and accuracy

In the detection of foot rot pathogens by the real-time PCR method of Fujiwara et al. (2021), a Ct value of <35 has been accepted as positive (NARO 2023; Tsukiori et al. 2023). Therefore, we calculated the detection performance (such as detection sensitivity and specificity) of this method when a Ct value <35 was defined as positive and a Ct value ≥35 as negative. We used 132 seedling tubers and 163 stem seedlings collected from actual fields as in the case of generation of ROC curves. The cross tables were created consisting of the binary values of positive or negative as the result of real-time PCR assay, and the binary values of whether the plant derived from each sample was actually diseased or healthy (Suppl. Fig.1). Each ability of detection performance was calculated as follows.

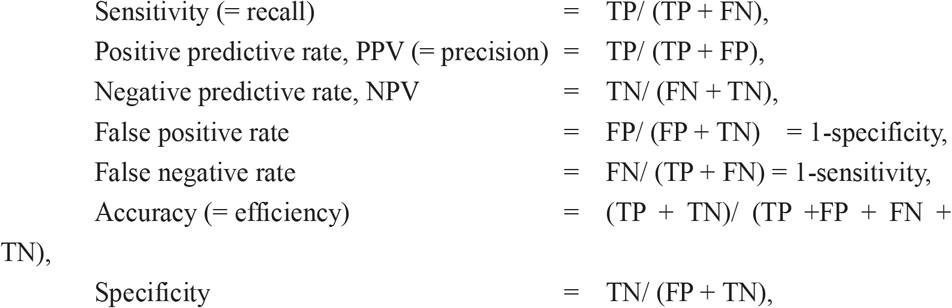

where TP; true positive, FN; false negative, TN; true negative, FP; false positive.

## Results

### The ROC curve of the real-time PCR using sweet potatoes collected from actual fields

The real-time PCR was performed on tubers and stems collected from fields where foot rot had been confirmed to occur, and Ct and Tm values were obtained for each sample (Suppl. Table 1 and 2). At this time, the corresponding main parts of tubers and stems of samples provided for the real-time PCR were cultivated for approximately three months, and the presence (“diseased”) or absence (“healthy”) of foot rot disease onset was also described in the table. To evaluate detection performance by the ROC curves, the results were organized into two cases: one where all Tm values, including abnormal ones, were considered (original case) and another where abnormal Tm values were excluded (Suppl. Table 3). The ROC curves were generated (Fig. 1), and the area under the ROC curve (AUC) was calculated for each case. AUC serves as an index of the ability of the real-time PCR results to correctly distinguish between diseased and healthy samples. A higher AUC value, approaching 1, indicates better detection accuracy. Additionally, when generating the ROC curves, detection accuracy at each Ct cutoff value was calculated, and the maximum accuracy for each case was determined (Suppl. Table 4). The ROC curves and these AUCs (0.7041571 - 0.9204286) indicated that the real-time PCR detection method developed by Fujiwara et al. (2021) had generally balanced detection performance for both tubers and stems in the field. Particularly, when stems were tested, the results were clearly distinguishable between samples infected with foot rot pathogen and healthy samples, and high AUCs (0.9204286 in original case and 0.9183741 in case in which abnormal Tm values were excluded) were obtained regardless of whether or not abnormal Tm values were excluded (Fig. 1c and d). On the other hand, testing on tubers resulted in a higher incidence of false positives at high Ct values (Suppl. Table 4). However, excluding abnormal Tm values improved the AUC from 0.7941571 (original case) to 0.8338827 (excluding abnormal Tm values) (Fig. 1a and b).

**Figure 1.**
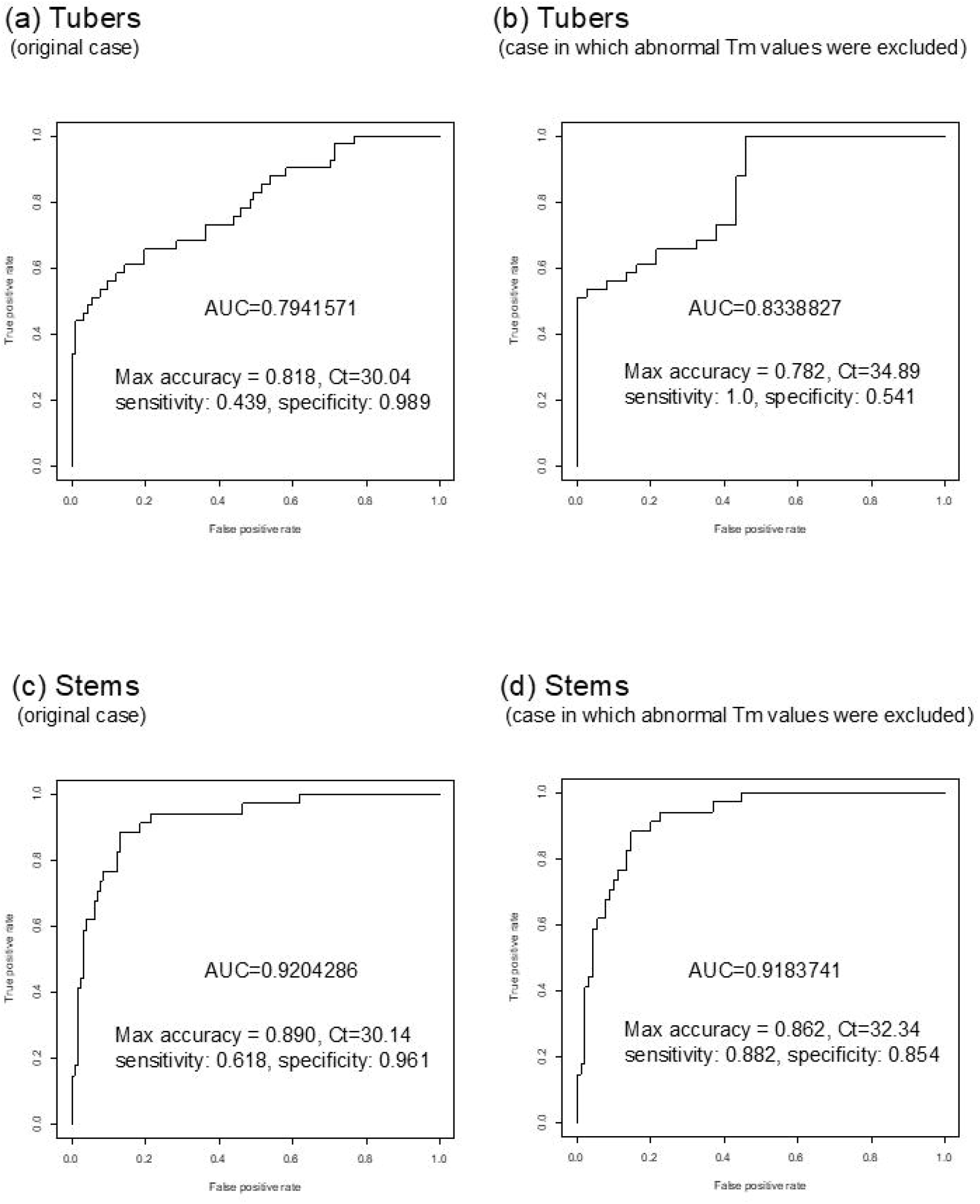
The receiver operating characteristic (ROC) curves of real-time PCR detection for foot rot disease with actual field samples. The ROC curves consisting of true positive rate and false positive rate were generated using the results of the real-time PCR and disease onset, when tubers and stems from areas where foot rot disease is widespread as samples. The ROC curves showed the true positive rate on the vertical axis and the false positive rate on the horizontal axis. AUC; area under the ROC curve, (a) original case of tubers, (b) case of tubers in which abnormal Tm values were excluded, (c) original case of stems, (d) case of stems in which abnormal Tm values were excluded.

### The detection performance of real-time PCR at Ct value 35 using sweet potatoes collected from actual fields

From the tables showing the results of real-time PCR and the onset of disease, we focused on minimizing false negatives with this detection method, and found that by setting a Ct value of 35 as the cutoff value, false negatives were less likely to occur in both tubers (sensitivity of 1.0) and stems (sensitivity of 0.9705882) (Suppl. Table 4). By setting a cutoff value of 35, the false negatives for the samples used in this study were 0 for tubers and 1 for stems. A false negative is when the detection results are negative despite the plant actually being infected with foot rot pathogen, and if the false negative rate is high, it can lead to serious errors in definitive diagnosis and health testing, especially. In other words, by using this method with a Ct value of 35 as the cutoff value, the healthiness of tubers and stems can be examined. Therefore, the detection performance at a Ct value of 35 was calculated. Detection performance, including sensitivity and specificity, was calculated by creating a cross-table according to the Materials and Methods section (Suppl. Fig. 1). The detection performance of each test for tubers and stems was examined by dividing the results into three cases: (1) distinguishing based on Ct values alone (Ct values < 35 or ≥ 35) without considering the Tm value (Fig. 2a and d) (original case), (2) distinguishing based on Ct values while excluding results with abnormal Tm values (Fig. 2b and e), and (3) distinguishing based on Ct values while interpreting results with abnormal Tm values as PCR negative (Fig. 2c and f).

**Figure 2.**
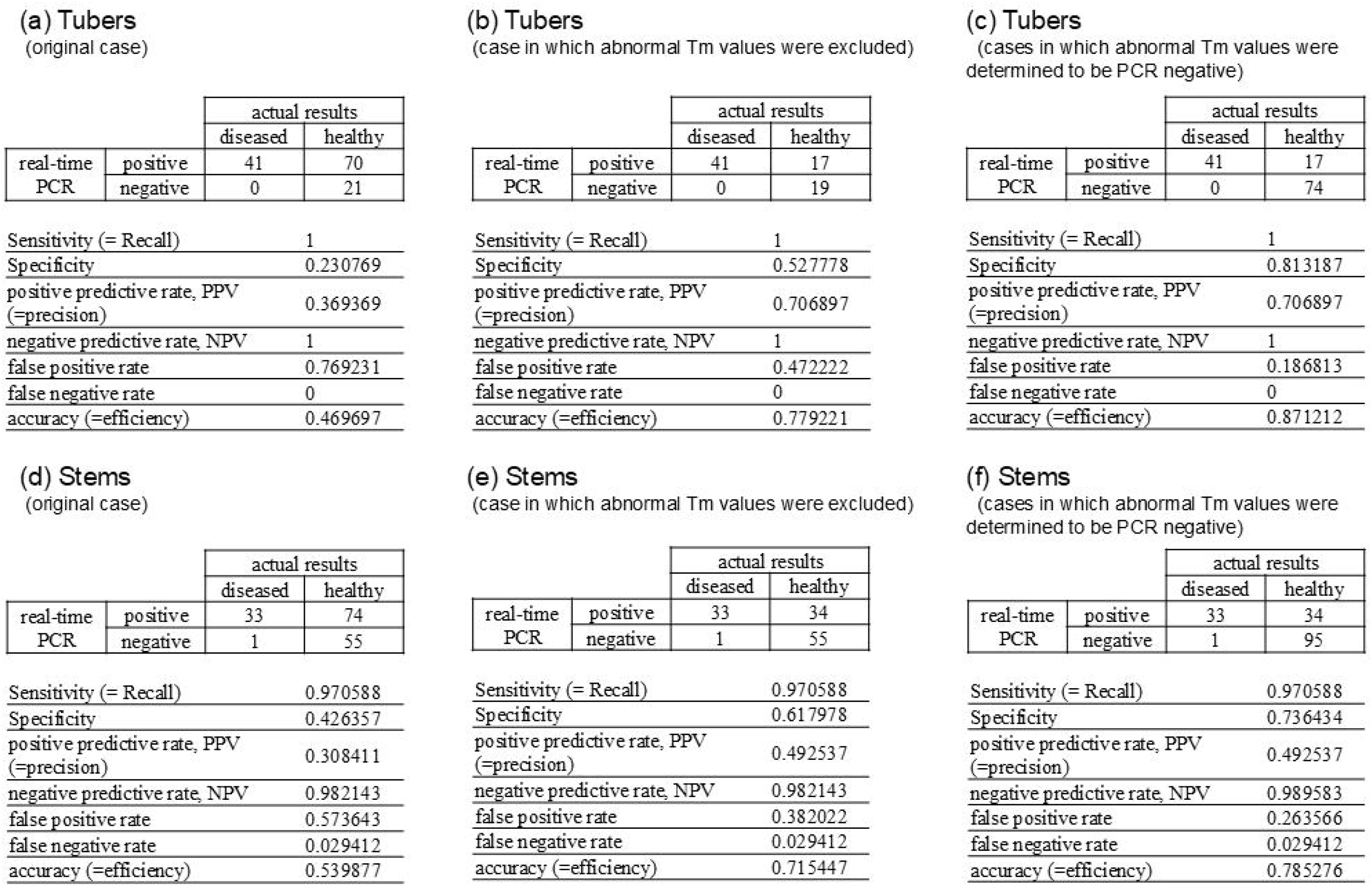
Detection performance of each test at a Ct value of 35. Using tubers and stems from areas where foot rot is prevalent as samples, we investigated the detection performance using real-time PCR results and disease onset, setting a cutoff Ct value of 35. We also confirmed cases in which abnormal Tm values were excluded and cases in which abnormal Tm values were determined to be PCR negative. (a) original case of tubers, (b) case of tubers in which abnormal Tm values were excluded, (c) case of tubers in which abnormal Tm values were determined to be PCR negative, (d) original case of stems, (e) case of stems in which abnormal Tm values were excluded, (f) case of stems in which abnormal Tm values were determined to be PCR negative.

For both tubers and stems, the accuracy (efficiency) was low when the Tm value was not considered in the analysis (original case), with values of 0.469697 for tubers and 0.539877 for stems. However, accuracy improved when abnormal Tm values were excluded and recalculated with values of 0.779221 for tubers and 0.715447 for stems, and high accuracy was achieved when all results showing abnormal Tm values were considered negative, reaching 0.871212 for tubers and 0.785276 for stems. Thus, the way in which abnormal Tm values were handled had a significant impact on specificity and related performance. Comparing these cases, the results indicated that highly accurate detection is possible by determining that a sample is PCR negative when the obtained Ct value is less than 35 but the Tm value is unexpected (Fig. 2).

## Discussion

Detection techniques, including PCR, have been widely developed in the field of plant protection, and there are many on-site diagnosis, identification, and testing methods that use these techniques. However, most of the detection techniques reported in research papers have few examples of field verification, and the actual detection performance is often unclear. In this study, we were able to evaluate the detection performance of our previously developed real-time PCR for foot rot detection using the contaminated tubers and stems in the field. When evaluating the detection performance using artificially contaminated plants, it is often difficult to determine whether the results match reality because the actual degree and amount of contamination is unknown. However, by using naturally contaminated plants, it is possible to evaluate the detection performance that is closer to on-site conditions. We had examined and secured a large number of contaminated tubers and stems at a time when the damage from foot rot disease had been particularly severe in Japan, and were therefore able to use them for this evaluation test.

In addition, in the actual fields where contaminated tubers and stems were collected, progress is being made in controlling foot rot disease, and the incidence of the disease is being suppressed. In this study, the evaluation of detection performance demonstrated that the real-time PCR method (Fujiwara et al. 2021) is effective in distinguishing between contaminated and healthy plants. In particular, it became clear that accuracy could be improved by “determining abnormal Tm values as PCR negative”, a setting that Fujiwara et al. had established during development. Fujiwara et al. (2021) reported that no false positives were observed below a Ct value of 32, but did not mention the occurrence of false negatives. In contrast, Tsukiori et al. (2023) confirmed that by using a Ct value of 35 as the cutoff, sweet potato pathogens other than the foot rot pathogen and healthy plants were not mistakenly detected. Furthermore, the ROC curve analysis and detection performance in this study demonstrated that a Ct value of 35 as the cutoff effectively minimizes false negatives. Thus, setting a Ct cutoff value of 35 ensures that as few contaminated plants as possible are overlooked, making it suitable for assessing the health status of seed tubers and stem seedlings in seedling propagation. Conversely, lowering the Ct cutoff below 35 enables the detection of highly contaminated plants more actively.

It is important to note that in medical diagnostics and other testing scenarios, cutoff values are often adjusted based on the specific purpose of the test. Similarly, in foot rot disease testing, the Ct value of 35 is not an absolute threshold. As far as our group has evaluated the performance of the real-time PCR, we have been able to perform detection with high accuracy by setting the cutoff value at Ct 35. However, the selection of Ct and Tm thresholds should be carefully considered based on the testing objective, field conditions, and testing capabilities. Furthermore, these parameters should be explicitly stated to ensure objective and reproducible evaluations. This study not only evaluated the detection performance of the real-time PCR for foot rot pathogens but also contributes to providing information on how to investigate detection performance and on its significance in various real-time PCR genetic detection methods.

## Acknowledgements

We thank the members of the Institute for Plant Protection, NARO and the Kyushu Okinawa Agricultural Research Center, NARO for helpful discussions.

## Author contributions

TF, YI, YT, HI and KF conceived and designed the study. TF, HI, and KF performed the experiments. TF, YI, YT and KF contributed to the data analysis. TF, YI, YT, HI, and KF contributed to the manuscript writing.

## Ethical declarations

### Ethical approval

This article does not contain any studies with human participants or animals.

### Data availability statement

The authors confirm that all data supporting the findings of this study are available within the article and its supplementary materials.

