## Supplemental_Figure_1 for "Evaluation of real-time PCR performance for detecting *Diaporthe destruens* in sweet potatoes": 07 Suppl Fig1 Cross table.pptx

### Slide 1
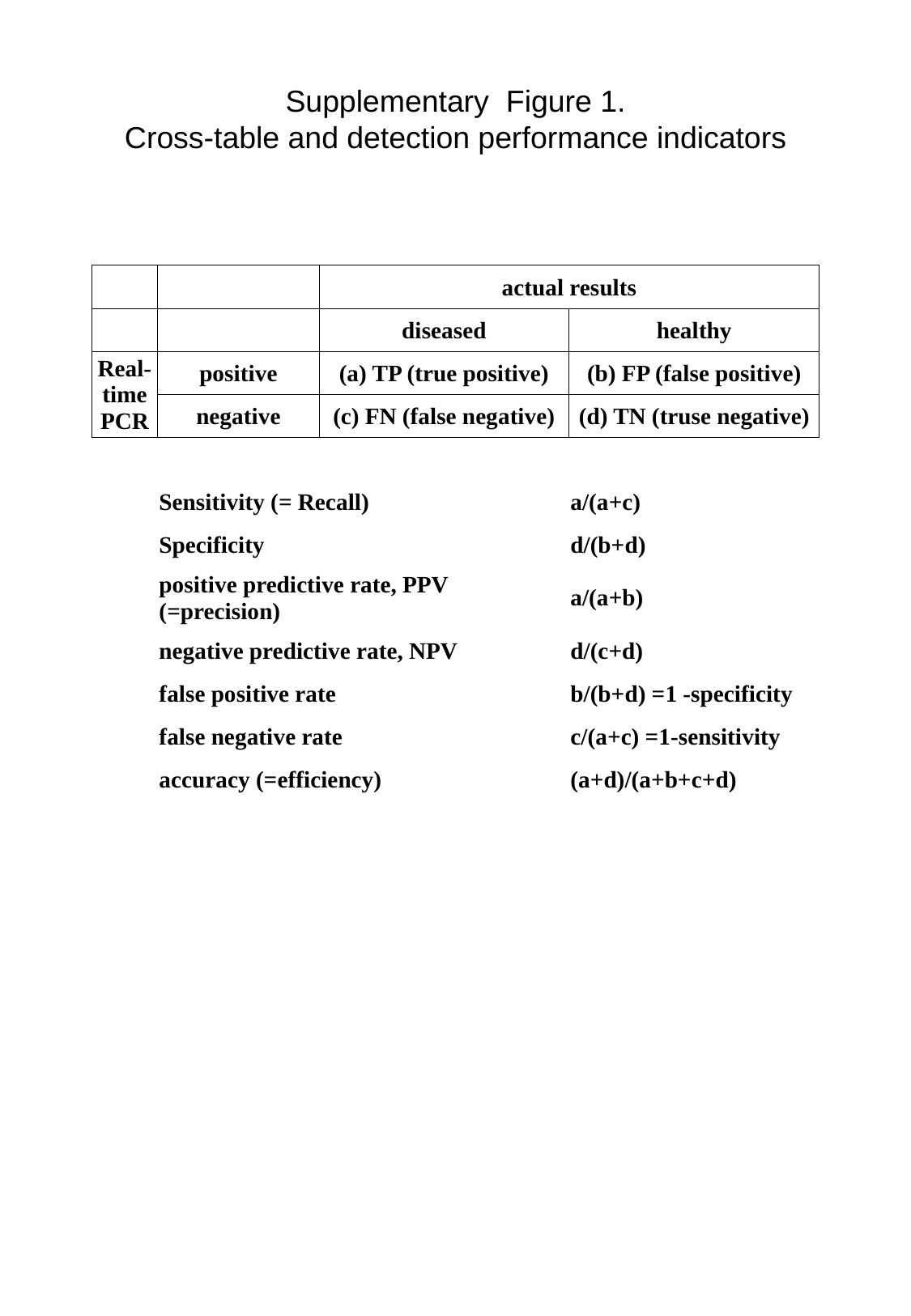

Supplementary Figure 1.
Cross-table and detection performance indicators
| | | actual results | |
| --- | --- | --- | --- |
| | | diseased | healthy |
| Real-time PCR | positive | (a) TP (true positive) | (b) FP (false positive) |
| | negative | (c) FN (false negative) | (d) TN (truse negative) |
| | Sensitivity (= Recall) | | a/(a+c) |
| | Specificity | | d/(b+d) |
| | positive predictive rate, PPV (=precision) | | a/(a+b) |
| | negative predictive rate, NPV | | d/(c+d) |
| | false positive rate | | b/(b+d) =1 -specificity |
| | false negative rate | | c/(a+c) =1-sensitivity |
| | accuracy (=efficiency) | | (a+d)/(a+b+c+d) |
