## Supplemental_Text_1 for "Evaluation of real-time PCR performance for detecting *Diaporthe destruens* in sweet potatoes": 08 Supplement Text 1. Example of ROC curve generation code.docx

**Supplemental Text 1. Example of ROC curve generation code in R**

rocdata <-read.table(“Table showing the relationship between Ct value and disease onset, eg., Suppl. Table 3 “)

rocdata

library(ROCR)

pred <- prediction(rocdata[,1], rocdata[,2])

perf <- performance(pred, "tpr", "fpr")

plot(perf)

png("roc-curve-stem.png") #Image output

plot(perf)

dev.off()

auc.tmp <- performance(pred,"auc") #Calculation of AUC

auc <- as.numeric(auc.tmp@y.values)

auc

table <- data.frame(Cutoff=unlist(pred@cutoffs),

TP=unlist(pred@tp), FP=unlist(pred@fp),

FN=unlist(pred@fn), TN=unlist(pred@tn),

Sensitivity=unlist(pred@tp)/(unlist(pred@tp)+unlist(pred@fn)),

Specificity=unlist(pred@tn)/(unlist(pred@fp)+unlist(pred@tn)),

Accuracy=((unlist(pred@tp)+unlist(pred@tn))/nrow(rocdata))

)

table　# Calculation of accuracy for each cutoff value

max(table$Accuracy) #Display of max accuracy
